# Dorsolateral septum GLP-1R neurons regulate feeding via lateral hypothalamic projections

**DOI:** 10.1101/2024.03.26.586855

**Authors:** Yi Lu, Le Wang, Fang Luo, Rohan Savani, Mark A. Rossi, Zhiping P. Pang

## Abstract

**Objective:** Although glucagon-like peptide 1 (GLP-1) is known to regulate feeding, the central mechanisms contributing to this function remain enigmatic. Here, we aim to test the role of neurons expressing GLP-1 receptors (GLP-1R) in the dorsolateral septum (dLS; dLS^GLP-1R^) and their downstream projections on food intake and determine the relationship with feeding regulation.

**Methods:** Using chemogenetic manipulations, we assessed how activation or inhibition of dLS^GLP-1R^ neurons affected food intake in *Glp1r-ires-Cre* mice. Then, we used channelrhodopsin-assisted circuit mapping, chemogenetics, and electrophysiological recordings to identify and assess the role of the pathway from dLS^GLP-1R^ neurons to the lateral hypothalamic area (LHA) in regulating food intake.

**Results:** Chemogenetic inhibition of dLS^GLP-1R^ neurons increases food intake. LHA is a major downstream target of dLS^GLP-1R^ neurons. The dLS^GLP-1R^→LHA projections are GABAergic, and chemogenetic inhibition of this pathway also promotes food intake. While chemogenetic activation of dLS^GLP-1R^→LHA projections modestly decreases food intake, optogenetic stimulation of the dLS^GLP-1R^→LHA projection terminals in the LHA rapidly suppressed feeding behavior. Finally, we demonstrate that the GLP-1R agonist, Exendin 4 enhances dLS^GLP-1R^ →LHA GABA release.

**Conclusions:** Together, these results demonstrate that dLS-GLP-1R neurons and the inhibitory pathway to LHA can regulate feeding behavior, which might serve as a potential therapeutic target for the treatment of eating disorders or obesity.

**Highlights:**

- Chemogenetic inhibition of dLS^GLP-1R^ neurons boosts food intake in mice
- dLS^GLP-1R^ neuron activation does not alter feeding, likely by collateral inhibition
- dLS^GLP-1R^ neurons project to LHA and release GABA
- Activation of dLS^GLP-1R^ →LHA axonal terminals suppresses food intake
- GLP-1R agonism enhances dLS^GLP-1R^ →LHA GABA release via a presynaptic mechanism

## 1. Introduction

Feeding behavior, a highly evolutionarily conserved process critical for survival, is orchestrated by multiple brain regions and related neural circuits in response to external cues and peripheral signals [1–3]. Abnormal activity within feeding circuits may lead to imbalanced food intake and energy expenditure, which can induce changes in body fat mass that can cause obesity and metabolism disease [1]. However, the neural circuit mechanisms controlling feeding behavior and how their dysfunction leads to disease remain poorly characterized.

Glucagon-like peptide 1 (GLP-1) is an incretin hormone and a post-translational cleavage product of preproglucagon encoded by the proglucagon *Gcg* gene, which is primarily produced in the L-cells of the intestine and a subpopulation of hindbrain nucleus tractus solitarius neurons [4–7]. GLP-1 actions are mediated by the GLP-1 receptor (GLP-1R), a class B subfamily G protein couple receptor, widely expressed in central and peripheral nervous systems [8–10]. Emerging evidence shows that endogenous GLP-1 signaling in both central and peripheral organs can regulate energy homeostasis, including food intake and glucose metabolism [11]. GLP-1 analogs have been used to treat type 2 diabetes [12], and GLP-1 receptor (GLP-1R) agonists, such as liraglutide, have recently been approved to treat obesity [11]. Although GLP-1R is widely expressed in brain, the physiological role of GLP-1R neurons in mediating food intake and body weight remains elusive.

One brain region that strongly expresses GLP-1R is the dorsolateral septum (dLS) [13; 14], a subcortical forebrain area that exerts influences over feeding behavior [15–18], sleep [19], learning and memory [20–22], mood and motivation [23; 24]. Previous studies have suggested GLP-1 acts in the lateral septum to control feeding. For example, infusions of GLP-1 into lateral septum can reduce food intake [25]; however, the involved cell types and neural circuits remain uncharacterized. The LHA has historically been considered a feeding control center that can serve to promote food seeking and intake [26; 27]. Emerging evidence showed that the lateral hypothalamic area (LHA) contains functionally opposing neuronal populations that induce diverse feeding behaviors [28; 29]. The projections from the septal area to the LHA are implicated in feeding behavioral regulation [30; 31], but whether dLS^GLP-1R^ neurons are involved in this pathway remains unknown.

In this study, we sought to determine the role of dLS^GLP-1R^ neurons in the regulation of food intake and to identify downstream targets mediating the effects. We found that chemogenetic inhibition of dLS^GLP-1R^ neurons increased food intake whereas chemogenetic activation of these neurons had no significant effect on feeding, likely due to local inhibition within dLS. Using channelrhodopsin-assisted circuit mapping, we show that a subset of dLS^GLP-1R^ neurons provide functional inhibitory connections onto neurons in LHA. Acute manipulations of this pathway bidirectionally modulate food intake. Collectively, our data demonstrate the role of dLS^GLP-1R^ neurons and their projections to LHA on septo-hypothalamic appetite regulation, providing insights into the central control of feeding behavior.

## 2. Method & Materials

### 2.1 Animals

All procedures involving mice were approved by the Rutgers University Institutional Animal Care and Use Committee and performed in accordance with the National Institute of Health guidelines for the care and use of laboratory animals. Adult (6-8 weeks old) *Glp1r-ires-Cre* and *Glp1r-ires-Cre:Ai14* male mice were used. Animals were bred in our facility and were maintained at constant temperature (∼22 °C), humidity (35%–55%), and circadian rhythm (12 h light/dark cycles). Food and water were available in their home cages *ad libitum*.

### 2.2. Stereotaxic surgery and AAV injections

The AAV-viruses used in this study include: pAAV-hSyn-DIO-EYFP (Catalog #27056-AAV9), retro-pAAV-Ef1a-DIO-EYFP (Catalog #27056-AAVrg), pAAV-hSyn-fDIO-hM4D(Gi)-mCherry-WPREpA (Catalog #154867-AAV8), pAAV-hSyn-fDIO-hM3D(Gq)-mCherry-WPREpA (Catalog #154868-AAV8), pAAV-Ef1a-fDIO-mCherry (Catalog #114471-AAV8), retro-AAV pEF1a-DIO-FLPo-WPRE-hGHpA (Catalog #87306-AAVrg), pAAV-EF1a-double floxed-hChR2(H134R)-EYFP-WPRE-HGHpA (Catalog #20298-AAV5), AAV-hSyn-DIO-hM4Di-mCherry (Catalog #44362-AAV9), AAV-hSyn-DIO-hM3Dq-mCherry (Catalog #44361-AAV9) (purchased from Addgene). Virus expression was allowed for a period of 3 weeks prior to experimental manipulation. For recording CNO-mediated effects of dLS^GLP-1R^ neurons, *Glp1r-ires-Cre* mice were injected bilaterally with either AAV-hSyn-DIO-hM4Di-mCherry or AAV-hSyn-DIO-hM3Dq-mCherry in the dLS allowing identification of dLS^GLP-1R^ neurons for electrophysiological recordings. For Channelrhodopsin-2 (ChR2)-assisted circuit mapping, *Glp1r-ires-Cre* animals were injected with pAAV-EF1a-double floxed-hChR2(H134R)-EYFP into dLS. The injection speed was 1 nL/s while the injection coordinates for dLS were: Anterior-Posterior (AP): +0.5 mm from bregma; Medial-Lateral (ML): ± 0.45 mm; Dorsal-Ventral (DV): -2.7 mm. The injection coordinates for the LHA were: AP: -1.2 mm from bregma; ML: ±1 mm; DV: -5.1 mm. Injection sites were confirmed by post hoc histological examinations in all animals reported in this study. 0.1-0.2 μl virus was used for the injection of AAV virus.

### 2.3. Whole-cell patch clamp electrophysiology in brain slices

Mice were deeply anesthetized with Euthasol and brains were quickly removed into ice-cold oxygenated cutting solution (in mM): 50 sucrose, 2.5 KCl, 0.625 CaCl_2_, 1.2MgCl_2_, 1.25NaH_2_PO_4_, 25 NaHCO_3_, and 2.5 glucose saturated with 95% O_2_/5% CO_2_. Coronal hypothalamic or dLS slices at 300μm were prepared using a vibratome (VT 1200 S; Leica). After 1 h recovery (33 °C) in artificial cerebrospinal fluid (ACSF) (in mM): 125 NaCl, 2.5 KCl, 2.5 CaCl_2_, 1.2 MgCl_2_, 1.25 NaH_2_PO_4_, 25NaHCO_3_, and 2.5 glucose. Slices were then transferred to a recording chamber and perfused with ACSF recording solution at 30 °C. Patch pipettes (6-9 MΩ) were pulled from borosilicate glass. For current clamp recordings, internal solution contains (in mM): 126 K-gluconate, 4 KCl, 10 HEPES, 4 Mg-ATP, 0.3 Na 2 -GTP, 10 phosphocreatine (pH to 7.2 with KOH); for voltage clamp synaptic recordings, internal solution containing (in mM): 40 CsCl, 90 K-Gluconate, 10 HEPES, 0.05 EGTA, 1.8 NaCl, 3.5 KCl, 1.7 MgCl 2, 2 Mg-ATP, 0.4 Na_4_-GTP, 10 Phosphocreatine and additional 5mM QX-314 (pH to 7.2 with CsOH). Whole-cell patch-clamp recordings were performed using an Axon 700B amplifier. Data were filtered at 2 kHz, digitized at 10 kHz and collected using Clampex 10.5 (Molecular Devices).

For chemogenetics recordings, 10 μM Clozapine-N-oxide (CNO) was applied to the bath solution. Neurons exhibiting an increase or decrease in resting membrane potential exceeding 5mV were considered significantly responsive. For photostimulation (470 nm) -evoked inhibitory post-synaptic currents (IPSCs) recordings, cells were held at -70mV in the presence of CNQX (20 μM).

### 2.4 Optical fiber placement

Bilateral optic fibers (RWD, 200 μm diameter core) were implanted into the LHA and fixed to the skull with cement. The coordinates used for the LHA were: AP: -1.2; ML: ±1; DV: -5.1 and AP: - 1.2; ML: ±1; DV: -5.0 for the optic fibers. After surgery, mice were housed individually and were allowed to recover for at least 3 weeks before conducting behavioral experiments. The optic fiber location was confirmed by post hoc histological examinations. Data were included only from animals with proper placement.

### 2.5 Chemogenetics and Food Intake

Food intake studies on chow were performed as previously described [32]. All animals were singly housed for 3 weeks following surgery and handled for 5 consecutive days before the assay to reduce stress responses. All studies were conducted in a homecage environment with *ad libitum* chow access. A trial consisted of assessing food intake from this study subjects after they received injections of saline or CNO (1mg/kg i.p.). Saline control experiments were performed with 1 day washout between conditions. CNO tests were conducted with 6 days washout between experiments to ensure the effects of CNO-mediated stimulation had dissipated. Mice with misplaced injections or poor expression were excluded from analysis after post hoc histological examination of mCherry expression. The investigators were blinded to all food intake measurements.

Dark-cycle hM3Dq/hM4Di feeding studies were conducted between 6:00pm to 9:00pm (lights off at 6:00 pm) and food intake was monitored 0.5 h, 1 h, 2 h and 3 h starting 30 min after i.p. injection. For post-fast refeed hM3Dq/hM4Di studies, animals were fasted overnight starting at 6:00pm and food was returned the following morning at 9:00am. Food intake was monitored 0.5 h, 1 h, 2 h and 3 h after i.p. injection. Light-cycle hM3Dq/hM4Di feeding studies were conducted in *ad libitum* fed mice between 9:00am to 12:00pm (lights on at 6:00 am) and food intake was monitored 0.5 h, 1 h, 2 h and 3 h after i.p. injection.

### 2.6. Optogenetics stimulation and food intake

Optical fibers (200 μm diameter core), NA 0.37 (Thorlabs) were connected the LED generator (473 nm for stimulation, Doric LEDFLP_4ch_1000). A terminal fiber was coupled to a 1.25 OD zirconium ferrule and a mating sleeve which allowed delivery of light to the brain. For in vivo photo-stimulation experiments, 10 ms pulses 1.8mW 473 nm blue light were given as 25 pulses/s every 1.5 s for 20 min. For the stimulation experiment, animals were provided only 0.5g food overnight to induce modest food deprivation. All mice were trained for 4 days to acclimate to the feeding schedule before testing. Food intake was measured at 20-, 40-, 60-min after optogenetic stimulation.

### 2.7. Open field test

Mice were introduced to the middle of a custom-made 45 x 45 cm square open field arena for 10 minutes. The mouse’s position was tracked from above with a video camera. Mouse position and time spent in the center/peripheral zones were analyzed with custom Python code. The mouse’s starting position is defined by the experimenter. Subsequent position is then automatically tracked in each frame by detecting the largest contour and calculating its center of mass. Metrics including the overall distance traversed, duration spent in central versus peripheral areas, and the count of entries into the central zone were calculated. An entry into the center zone was defined as when the center of mass of the animal image enters the center square. Distance traveled and time in center zone were calculated with the custom script. Investigators were blinded to the condition during analysis.

### 2.8. Light-dark box test

The light-dark box consisting of one dark compartment (6 x 5 x 7 cm) and one illuminated compartment (6 x 5 x 7 cm) connected by a door. At the beginning of the experiments, the animals were individually placed into the light compartment facing the dark compartment and then were allowed to freely explore for 5 min. Time spent in the light zone and number of entries into the light zone were measured using SmartCage tracking system (AfaSci).

### 2.9. Elevated Plus Maze test

The elevated plus maze consisted of a plus-shaped platform with four intersecting arms: two open arms (30 cm × 5 cm, wall-free) and two enclosed arms (30 cm × 5 cm) surrounded by 15-cm-high walls. The maze was elevated 55 cm from the ground. Mice were placed in the center of the apparatus facing an open arm and then allowed to freely explore the maze for 10 min. Light in the open arms was kept at 40 lux. The time spent in the open arm was recorded using a custom Python script designed to automatically detect and track the movement of the animal. Initially, the script captured video input from a USB camera, and the user defined the region of interest (ROI) for both arms of the EPM. The script then identifies the animal’s position in each frame by detecting the largest contour and calculating its center of mass. These data were used to track the animal’s movement, recording coordinates at each time point. The script then calculated the duration spent in both the open and closed arms along with the number of entries into each arm. An entry was defined as when the center of mass of the animal enters an arm.

### 2.10. Statistical analysis

Statistical analyses were performed using GraphPad Prism 9. Data were analyzed using Student’s two-tailed t-tests for comparisons between two groups and one-way ANOVA followed by Dunnett’s multiple comparisons tests or two-way ANOVA followed by Sidak’s multiple comparisons tests for comparisons between 3 or more groups. Data are presented as mean ±SEM. A p-value < 0.05 was considered statistically significant.

## 3. Results

### 3.1. Chemogenetic Inhibition of dLS^GLP-1R^ increases food intake

To examine the role of dLS^GLP-1R^ neurons on food intake, we injected pAAV-hSyn-fDIO-hM4D(Gi)-mCherry in the dLS region of *Glp1r-ires-Cre* mice to express the inhibitory designer receptors exclusively activated by designer drugs (DREADDs)-hM4Di in dLS^GLP-1R^ neurons (Figure 1A&B). Ex-vivo slice whole-cell recordings confirmed that CNO potently inhibited dLS^GLP-1R^ neurons transduced with the hM4Di. The basal firing rate was reduced, and membrane potentials became more hyperpolarized following CNO application in acute brain slices (Figure 1C). To establish whether these neurons regulate food intake, we measured food intake after chemogenetic inhibition with intraperitoneal (i.p.) administration of CNO (1mg/kg). We first measured ad libitum food intake during dark and light cycles and found that food intake was significantly increased after chemogenetic inhibition (Figure 1 D&E). We also tested this after fasting. Again, we observed that animals with chemogenetic inhibition of dLS^GLP-1R^ neurons ate more food than controls (Figure 1F). These experiments indicate that dLS^GLP-1R^ neurons are involved in food intake regulation.

**Figure 1.**
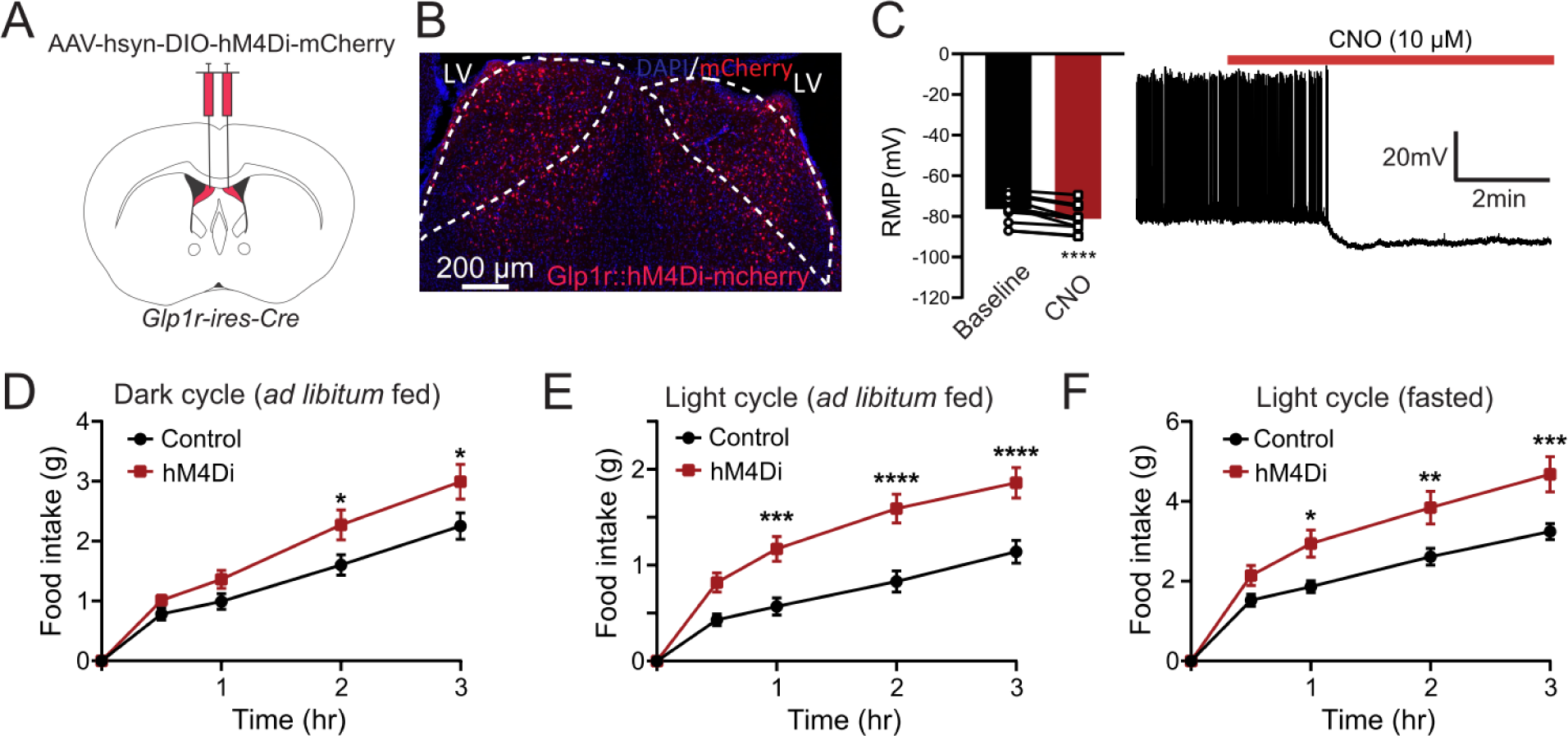
Inhibition of dLS^GLP-1R^ neurons acutely increases food intake. (A) Brain schematic of viral injection for chemogenetic dLS^GLP-1R^ neurons inhibition. (B) Transduction of dLS^GLP-1R^ neurons with hM4Di-mCherry. (C) CNO application hyperpolarized the resting membrane potential and reduced the firing rate of dLS^GLP-1R^ neurons (paired t-test, t(12)=6.281, p<0.0001, n=12 cells from 3 mice). (D-F) Chemogenetic inhibition of dLS^GLP-1R^ neurons increased food intake (D) during the dark cycle when fed (two-way ANOVA, main effect of Group: F(1,120)=14.44,p=0.0002; main effect of time: F(4,120)=71.72, p<0.0001, no interaction between Group and Time: F (4,120)=1.691, p=0.1566), (E) during the light cycle when fed (two-way ANOVA, main effect of Group: F(1,120)=53.89, p<0.0001; main effect of time: F(4,120)=58.53, p=<0.0001, interaction between Group and Time: F(4,120)=4.284, p=0.0028) and (F) refeeding following an overnight fast (two-way ANOVA, main effect of Group: F(1,120)=28.55, p<0.0001; main effect of Time: F(4,120)=68.25, p<0.0001, interaction between Group and Time: F(4,120)=2.463, p=0.0488). n=13 mice per group. Data are presented as mean ± SEM. Sidak’s multiple comparisons test: *p<0.05; **p<0.01; ***p<0.001, ****p<0.0001.

### 3.2. Activation of dLS^GLP-1R^ neurons has no significant effect on food intake

Because chemogenetic inhibition of dLS^GLP-1R^ neurons increased food intake, we hypothesized that activation of these neurons would conversely suppress feeding. To test this, we expressed stimulatory Gq-coupled hM3Dq-DREADD in dLS^GLP-1R^ neurons (Figures 2A&B). In contrast to chemogenetic inhibition (Figure 1C), CNO application failed to induce consistent effects on membrane properties in dLS^GLP-1R^ neurons (Figure 2C). After CNO application, 35% (6/17) of neurons showed a hyperpolarized membrane potential and reduced firing rate, whereas 29% (5/17) of neurons showed a depolarized membrane potential and increased firing rate. We then, measured food intake after CNO in vivo. Consistent with the varied in vitro recording responses, we also did not observe significant effects of CNO administration on food intake in any of the feeding tests (Figure 2D-F). This contradicted our initial hypothesis that activation of dLS^GLP-1R^ neurons would suppress food intake.

**Figure 2.**
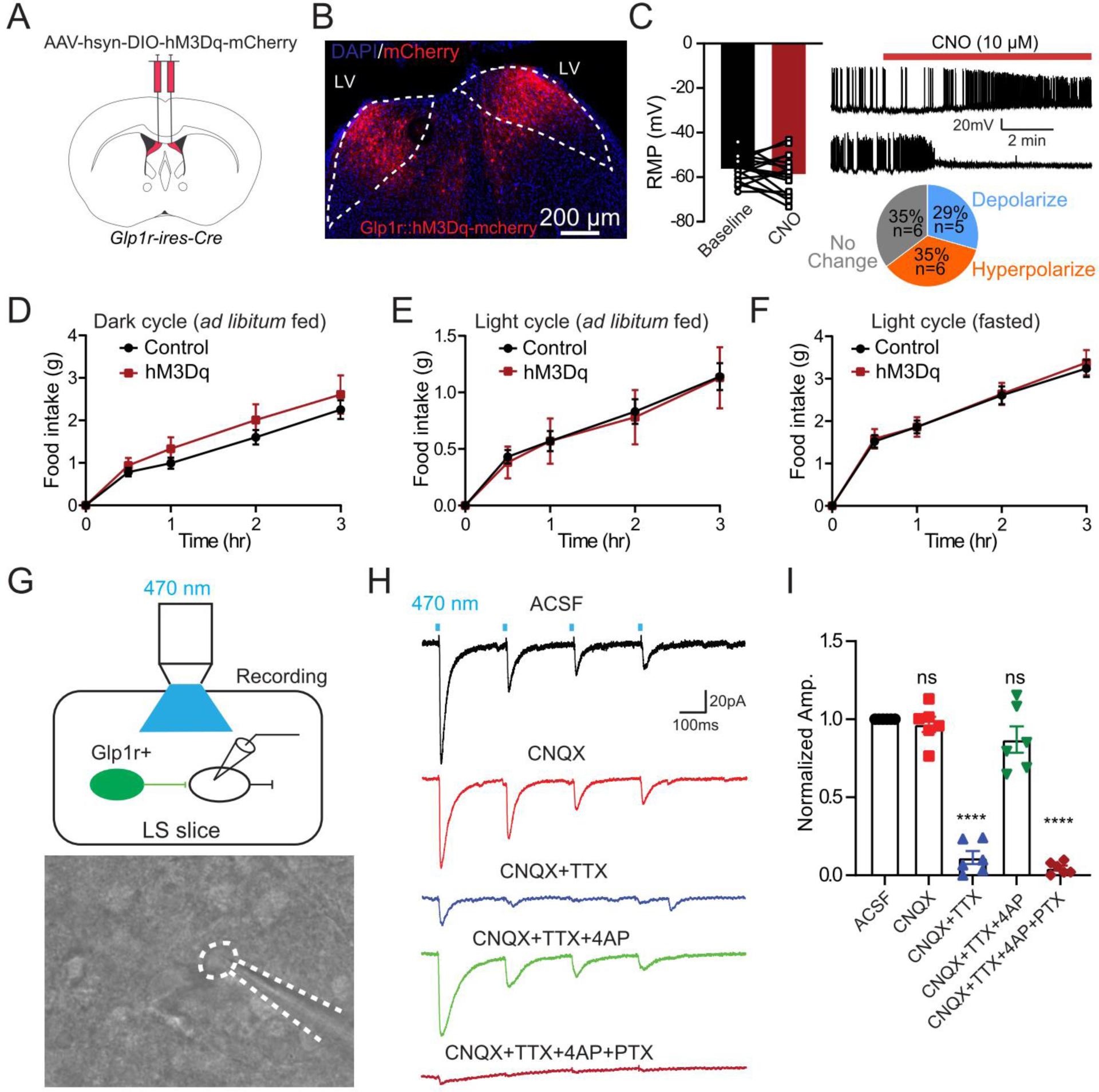
Activation of dLS^GLP-1R^ neurons has no significant effect on food intake. (A) Brain schematic of viral injection for dLS^GLP-1R^ neurons activation. (B) Transduction of dLS^GLP-1R^ neurons with hM3Dq-mCherry. (C) CNO application has no significant effect on the resting membrane potential and the firing rate of dLS^GLP-1R^ neurons (paired t-test, t(17), p=0.2523, n=17 cells from 3 mice), after CNO application, 35% of neurons exhibited resting membrane potential hyperpolarization, and 29% of neurons exhibited resting membrane potential depolarization. (D-F) Chemogenetic activation of dLS^GLP-1R^ neurons has no significant effect on food intake (D) during the dark cycle when fed (two-way ANOVA, no main effect of Group: F(1,105)=3.402, p=0.0679; main effect of Time: F(4,105)=36.10, p<0.0001, no interaction between Group and Time: F(4,105)=0.3067, p=0.8730.), (E) during the light cycle when fed (two-way ANOVA, no main effect of Group: F(1,105)=0.06226, p=0.8034; main effect of Time: F(4,105)=18.67, p<0.0001, no interaction between Group and Time: F(4,105)=0.01724, p=0.9994), and (F) refeeding following an overnight fast (two-way ANOVA, no main effect of Group: F(1,105)=0.1432, p=0.7059; main effect of Time: F(4,105)=84.5, p<0.0001, no interaction between Group and Time: F (4,105)=0.04575, p=0.9960). n=13 control and 10 hM3Dq mice. (G-I) Local inhibitory connections in dLS. Schematic showing the experiment used to record postsynaptic currents in local dLS neurons induced by optogenetic stimulation of dLS^GLP-1R^ neurons (G). In brain slice recordings, blue light stimulation evoked robust IPSCs, which were blocked by picrotoxin but not CNQX (H-I) (one-way ANOVA, F (4,25)=98.81, p<0.0001, ****Sidak’s multiple comparisons test p<0.0001 vs. ACSF). n=6 cells from 3 mice. Data are presented as mean ± SEM.

We previously demonstrated strong collateral inhibition among dLS neurons [21]. We thus hypothesized that collateral inhibition within dLS may prevent chemogenetic activation from influencing food intake. To directly test this possibility, we injected Cre-dependent AAVs expressing channelrhodopsin-2 (ChR2) with EYFP into dLS of *Glp1r-ires-Cre* mice and recorded post-synaptic responses from nearby EYFP-negative neurons (Figure 2G). Indeed, light stimulation of ChR2-expressing dLS^GLP-1R^ neurons evoked robust picrotoxin (PTX)-sensitive IPSCs in some neighboring EYFP-negative dLS neurons (6/11) (Figure 2H&I). Taken together, these findings suggest that chemogenetic activation of dLS^GLP-1R^ neurons has no significant effect on food intake, which is likely mediated by local inhibition in dLS.

### 3.3. dLS^GLP-1R^ neurons has no significant impact on stress related behavior

Because previous studies showed that alterations in basal anxiety levels or locomotion may affect feeding behavior [33; 34], we investigated whether dLS^GLP-1R^ neural regulation of food intake was secondary to the potential changes in anxiety-like behavior or locomotion. To test this possibility, we performed open-field test, light-dark box test and elevate plus maze test in *Glp1r-ires-Cre* mice transduced with the DREADD-hM3Dq or DREADD-hM4Di in dLS^GLP-1R^ neurons. We observed no changes in basal anxiety-like behavior during chemogenetic inactivation or activation of dLS^GLP-1R^ neurons in these behavioral tests (Figure S1), suggesting that altered feeding behavior was not induced by changes in basal anxiety levels.

### 3.4. dLS^GLP-1R^ neurons project to LHA

Because dLS^GLP-1R^ neuron inhibition increases food intake (Figure 1), we hypothesized that these neurons regulate feeding via long range projections. To determine the downstream targets of dLS^GLP-1R^ neurons, we injected *Glp1r-ires-Cre* mice in dLS with a Cre-dependent AAVs expressing EYFP to visualize axons innervating distal brain regions (Figure 3A). We observed EYFP+ soma in dLS (Figure 3B) and EYFP+ fibers in LHA (Figure 3C), suggesting that dLS^GLP-1R^ neurons send projections to LHA. To further confirm this pathway, we injected a retrograde tracer, retroAAV-Ef1a-DIO-EYFP, in LHA of *Glp1r-ires-Cre* mice crossed with a Cre-dependent tdTomato reporter line (Ai14) to visualize GLP-1R-expssing neurons (Figure 3D). Unsurprisingly, tdTomato^+^ cells were distributed evenly throughout the dLS, and EYFP retrogradely labeled dLS neurons were observed (Figure 3E&F). Overall, these experiments showed that dLS^GLP-1R^ neurons project to LHA. To further confirm this projection pathways and to reveal the properties of their neurotransmitter release, we performed electrophysiological recordings. In *Glp1r-ires-Cre* mice we injected Cre-dependent ChR2-EYP in the dLS and made whole cell patch clamp recordings from LHA neurons (Figure 3G). Reliable optogenetic-evoked IPSCs were recorded in LHA neurons (5/8, 62.5%). The blue light-induced PSCs were abolished upon application of TTX and recovered upon application of 4-AP, suggesting monosynaptic connectivity. Light-evoked PSCs were also abolished upon application of the GABA_A_ receptor antagonist picrotoxin (PTX), confirming that dLS^GLP-1R^→LHA neurons are GABAergic (Figure 3H&I). Taken together, these results show that dLS^GLP-1R^ neurons project to LHA where they form functional inhibitory synapses.

**Figure 3.**
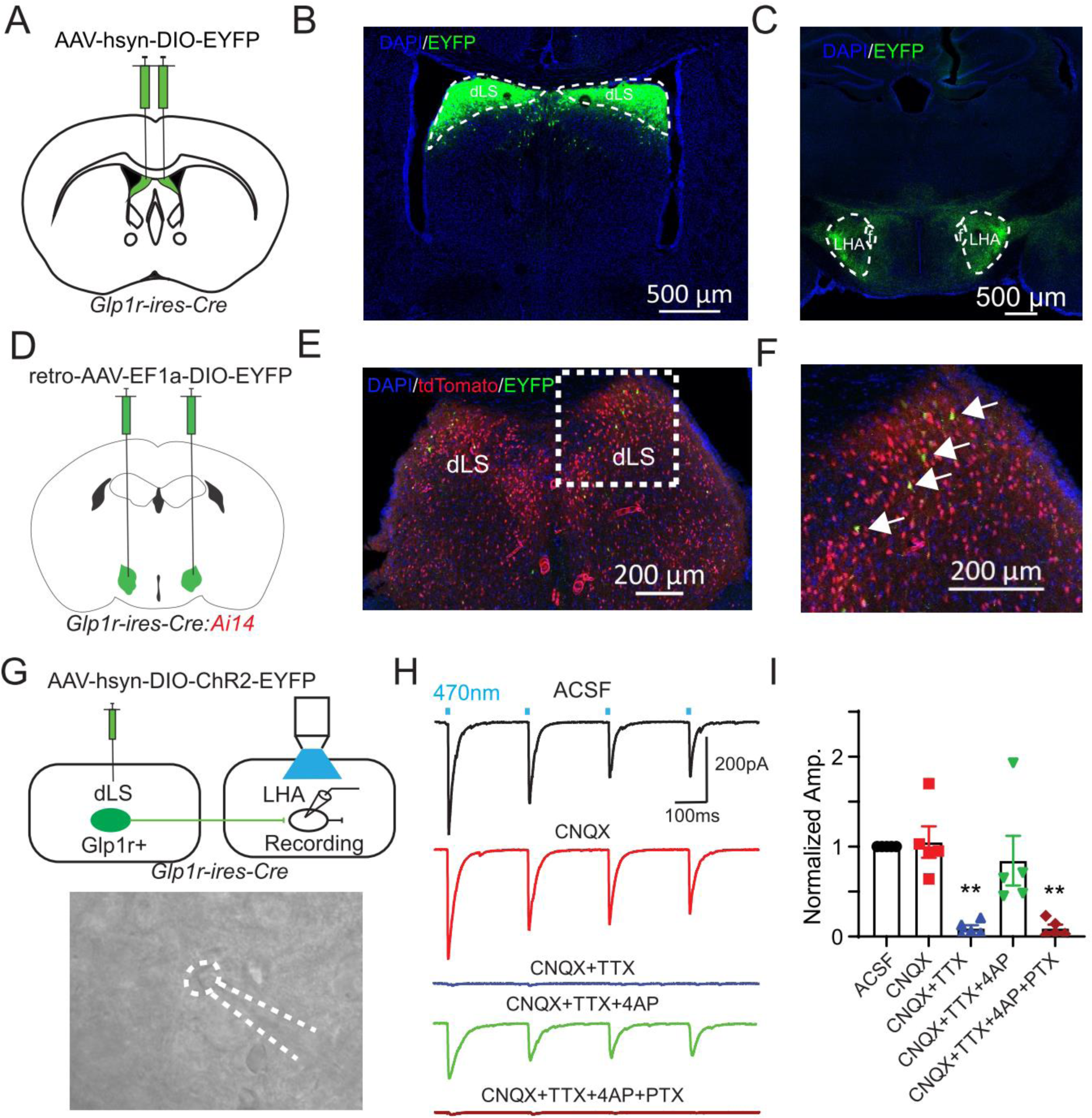
dLS^GLP-1R^ neurons project to LHA. (A) *Glp1r-ires-Cre* mice were injected in dLS with pAAV-hSyn-DIO-EYFP bilaterally. (B and C) Viral-mediated expression of EYFP in dLS^GLP-1R^ neurons soma (B) and axonal projections in the LHA (C) in *Glp1r-ires-Cre* mice. (D) *Glp1r-ires-Cre:Ai14* mice were injected in LHA with retro-pAAV-Ef1a-DIO-EYFP bilaterally. (E and F) Images of coronal brain sections containing the dLS. White arrows indicate GLP-1R+/EYFP+ neurons. (G) Schematic of experiment used to record postsynaptic currents in LHA neurons induced by optogenetic stimulation of dLS^GLP-1R^ neurons. *Glp1r-ires-Cre* mice were injected in dLS with pAAV-EF1a-double floxed-hChR2(H134R)-EYFP-WPRE-HGHpA bilaterally and LHA neurons were patched. (H) Representative trace of photostimulation (470 nm LED)-evoked IPSC in LHA neurons, which can be blocked by picrotoxin but not CNQX. (I) Normalized IPSC amplitude before and after CNQX, TTX, 4AP and PTX application (one-way ANOVA, F(4,20)=10.7, p<0.0001, **Sidak’s multiple comparisons test p<0.01 vs. ACSF). n=5 cells from 3 mice. Data are presented as mean ± SEM.

### 3.5. dLS^GLP-1R^ neurons regulate food intake via projections to LHA

We next asked if dLS^GLP-1R^→LHA manipulations are sufficient to influence food intake. We used chemogenetic and optogenetic approaches to manipulate dLS^GLP-1R^→LHA projections and probed their ability to regulate food intake. First, we injected *Glp1r-ires-Cre* mice with retroAAV-DIO-flp in LHA and AAV-fDIO-hM4Di-mCherry or AAV-fDIO-mCherry in dLS to express the inhibitory DREADDs only in the dLS^GLP-1R^ neurons that project to the LHA (Figure 4A&B). We confirmed the effectiveness of chemogenetic inhibition by in vitro electrophysiology (Figure 4C). Similar to global manipulations of dLS^GLP-1R^ neurons (Figure 1), we observed significant food intake augmentation in inhibiting dLS^GLP-1R^→LHA neurons (Fig. 4D-F), without affecting anxiety-like behaviors (Figure S2). These data suggest that LHA is likely one of the main targets of dLS^GLP-1R^ neurons that mediates food intake behavior.

**Figure 4.**
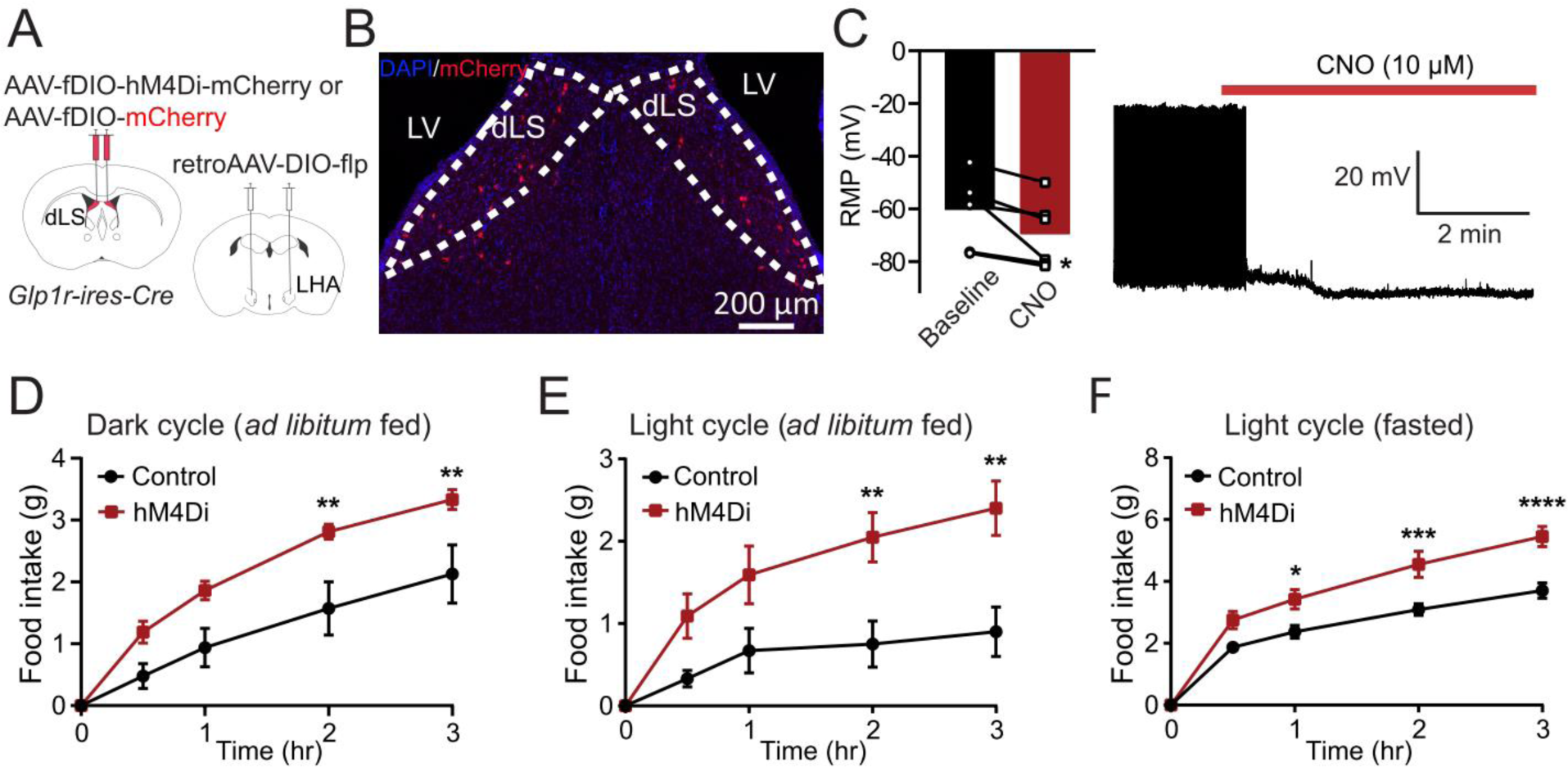
dLS^GLP-1R^→LHA inhibition increases feeding. (A) Brain schematic of viral injection for dLS^GLP-1R^→LHA neuron inhibition. (B) Transduction of dLS^GLP-1R^→LHA neurons with hM4Di-mCherry. (C) CNO application hyperpolarized the resting membrane potential and reduced the firing rate of dLS^GLP-1R^→LHA projection neurons (paired t-test, t(6)=2.897, p=0.0339, n=6 cells from 3 mice). (D-F) Chemogenetic inhibition of dLS^GLP-1R^→LHA projection neurons increased food intake (D) during the dark cycle when fed (two-way ANOVA, main effect of Group: F(1,40)=26.01, p<0.0001; main effect of Time: F(4,40)=36.65, p<0.0001, no interaction between Group and Time: F(4,40)=1.993, p=0.1142), (E) during the light cycle when fed (two-way ANOVA, main effect of Group: F(1,60)=27.88, p<0.0001; main effect of Time: F(4,60)=11.68, p<0.0001, no interaction between Group and Time: F(4,60)=2.344, p=0.0649), and (F) refeeding following an overnight fast (two-way ANOVA, main effect of Group: F(1,55)=44.88, p<0.0001; main effect of Time: F(4,55)=104, p<0.0001, interaction between Group and Time: F(4,55)=3.795, p=0.00085). n=7 control an d6 hM4Di mice. Sidak’s multiple comparisons test: *p<0.05, **p<0.01, ***p<0.001, ****p<0.0001. Data are presented as mean ± SEM.

Given the possibility that collateral inhibition affects the behavioral expression of activation of dLS^GLP-1R^ neurons (Figure 2), we hypothesized that restricting activation of dLS^GLP-1R^ neurons to those projecting to the LHA would suppress food intake. To test this hypothesis, we injected *Glp1r-ires-Cre* mice with retroAAV-DIO-flp in LHA and AAV-fDIO-hM3Dq-mCherry or AAV-fDIO-mCherry in dLS to express the excitatory DREADDs in dLS^GLP-1R^→LHA projections (Figure 5A&B). In contrast to the global dLS^GLP-1R^ activation, we observed consistent membrane depolarization of hM3Dq-expressing neurons when CNO was applied *ex vivo* (Figure 5C). Upon CNO administration *in vivo*, we observed no change in food intake in either the dark or light cycles when animals were *ad libitum* fed (Figures 5D&E). However, food intake was significantly reduced when animals were tested in the light cycle following fasting (Figure 5F). Anxiety-like behavior was again unchanged in these animals (Figure S3).

**Figure 5.**
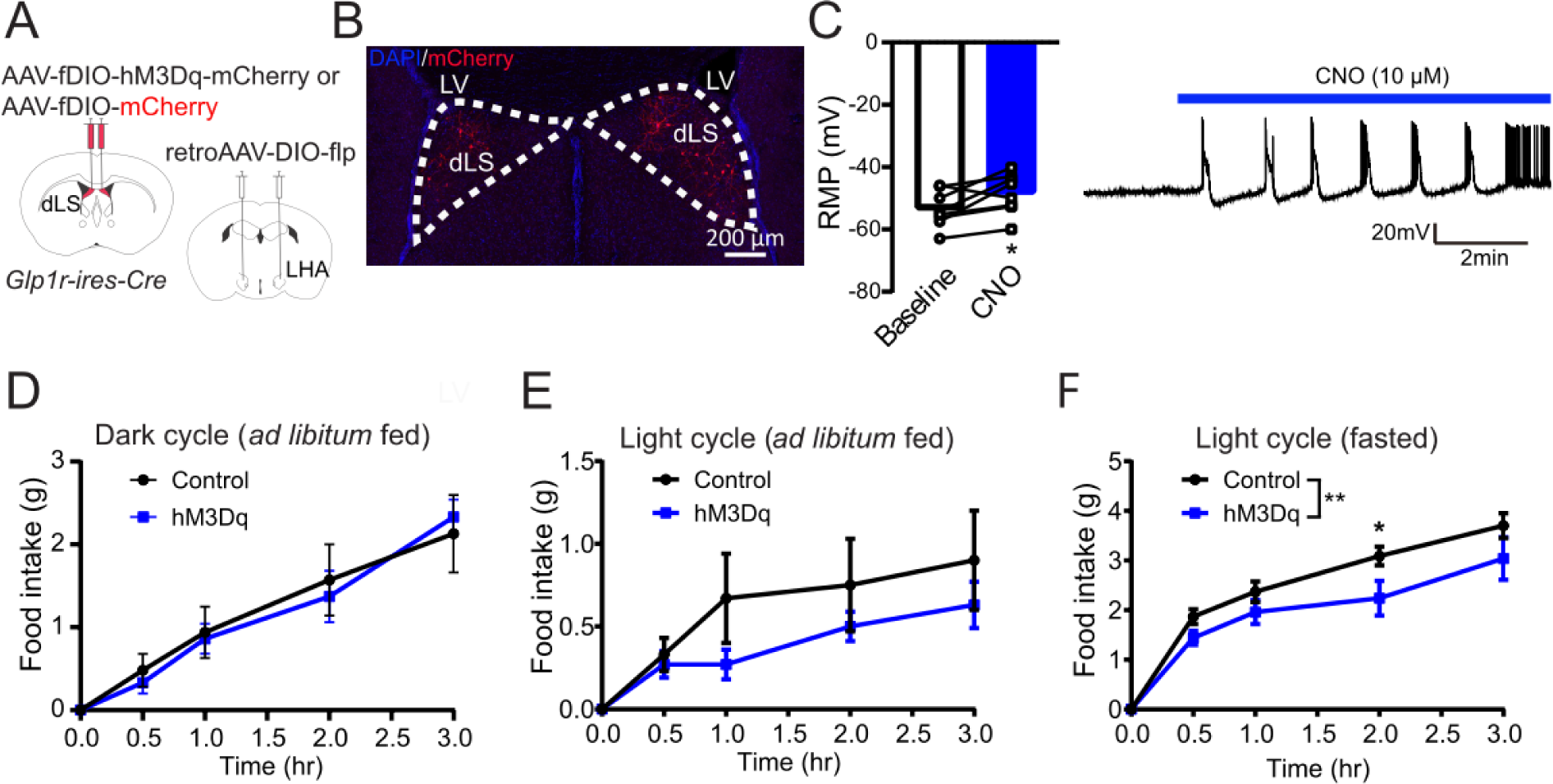
Activation of dLS^GLP-1R^ projection neurons has no significant effect on food intake. (A) Brain schematic of viral injection for dLS^GLP-1R^→LHA neuron activation. (B) Transduction of dLS^GLP-1R^→LHA neurons with hM3Dq-mCherry. (C) CNO application depolarized the resting membrane potential and increased the firing rate of dLS^GLP-1R^→LHA projection neurons (paired t-test, t(8)=2.702, p=0.0306, n=8 cells from 3 mice). (D-F) Chemogenetic activation of dLS^GLP-1R^→LHA projection neurons has no significant effect on food intake (D) during the dark cycle when fed (two-way ANOVA, no main effect of Group: F(1,45)=0.07858, p=0.7805; main effect of Time: F(4,45)=23.01, p<0.0001, no interaction between Group and Time: F(4,45)=0.1826, p=0.9463), (E) during the light cycle when fed (two-way ANOVA, a marginally significant main effect of Group: F(1,50)=3.272, p=0.0765; main effect of Time: F(4,50)=6.024, p=0.0005, no interaction between Group and Time: F(4,50)=0.4554, p=0.7680). (F) Chemogenetic activation suppressed food intake during refeeding following an overnight fast (two-way ANOVA, main effect of Group: F(1,50)=11.01, p=0.0017; main effect of Time: F(4,50)=64.4, p<0.0001, no interaction between Group and Time: F(4,50)=1.013, p=0.4096). n=7 control and 5 hM3Dq mice. Data are presented as mean ± SEM.

To further probe whether acute activation of the dLS^GLP-1R^→LHA pathway could suppress feeding, we next performed projection-specific optogenetic stimulation. We expressed Cre-dependent ChR2-EYFP in dLS of *Glp1r-ires-Cre* mice and implanted optical fibers in the LHA to selectively stimulate the dLS^GLP-1R^→LHA pathways (Figure 6A&B). Indeed, optogenetic activation of dLS^GLP-1R^→LHA nerve terminals significantly suppressed feeding in fasted mice, which returned to control levels when stimulation ceased (Figure 6C).

**Figure 6.**
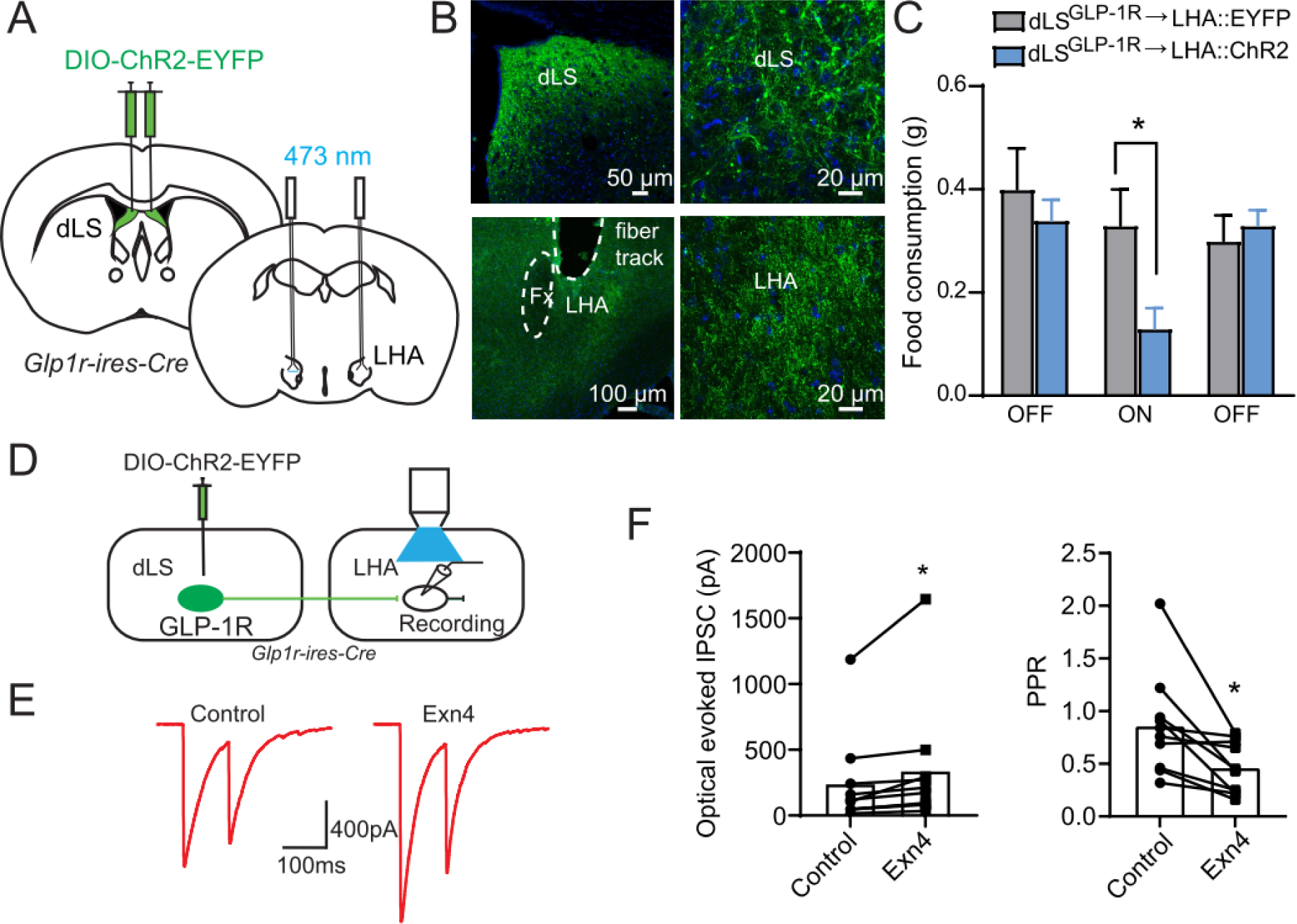
dLS^GLP-1R^→LHA activation suppresses feeding. (A and B) ChR2-EYFP expression in the dLS (A) and axonal projections in the LHA (B) in *Glp1r-ires-Cre* mice. (C) Photostimulation of dLS^GLP-1R^→LHA projections significantly suppressed food intake (two-way ANOVA, main effect of Group: F(1,36)=3.382, p=0.0742; main effect of Stimulation: F(2,36)=3.816, p=0.0314, no interaction between Group and Stimulation: F(2,36)=2.576, p=0.0900. *Sidak’s multiple comparisons test p<0.05). Data are presented as mean +SEM. n=6 controls and 8 ChR2 mice. (D) Experimental paradigm for recording postsynaptic currents in LHA slices induced by optogenetic stimulation of dLS^GLP-1R^ neurons. (E) Representative evoked IPSCs after paired-pulse optogenetic stimulation. (F) Quantification of evoked IPSCs (paired t-test, t(10)=2.312, p=0.0461) and PPR (paired t-test, t(10)=3.135, p=0.0120) before and after Ex4 application. n=10 cells from 3 mice. *p<0.05.

Finally, since these dLS^GLP-1R^→LHA neurons express GLP-1R, we asked if GLP-1R-mediated signaling affects dLS^GLP-1R^→LHA synaptic strength[35]. We recorded from LHA neurons that receive inhibitory synaptic input from dLS^GLP-1R^ neurons (Figure 6D). Consistent with our hypothesis, optically induced IPSCs in the LHA neurons were enhanced by application of the GLP-1R agonist, Exendin 4 (Exn-4) (Figure 6E&F). This synaptic augmentation is accompanied by a reduction of paired-pulse ratio (PPR), suggesting the involvement of a presynaptic mechanism.

Overall, these experiments establish that dLS^GLP-1R^ neurons and their inhibitory projections to the LHA regulate food intake independent of changes in basal anxiety levels.

## 4. Discussion

Our data demonstrate that dLS^GLP-1R^ neuron inhibition potentiates food intake without affecting changes in anxiety-like behavior and locomotion. Previous studies have shown that the projections from the lateral septum to thalamic nuclei mediate stress and anxiety-like behavior, including both anxiolytic and anxiogenic effects [36–41]. We found that manipulations of dLS^GLP-1R^ neurons or their projections to the LHA failed to affect anxiety-like behavior while still influencing food intake. This suggests that this population is selectively involved in the control of feeding behavior. However, the lack of obvious effects on anxiety-like behavior in this study may be due to the possibility that we manipulated neurons that mediate both anxiolytic and anxiogenic behaviors.

Consistent with our observations, emerging evidence has demonstrated that the lateral septum contributes to appetite suppression [17; 30] and food-seeking behaviors [31]. We extend these findings by demonstrating a selective role of dLS^GLP-1R^ neurons in regulating food intake. In the current study, we show that chemogenetic dLS^GLP-1R^ neuron inhibition increases food intake; however, chemogenetic activation of the same population failed to affect food intake. It has previously been shown that GABAergic neurons comprise more than 90% of all LS neurons [42]. We therefore hypothesized that local inhibitory connections within the dLS result in some cells being inhibited during dLS^GLP-1R^ neuron activation. Using optogenetics-assisted circuit mapping, we indeed recorded GABA release in dLS after GLP-1R neuron activation. In support of this, a previous study showed that activation of LS-A_2A_R (adenosine A_2A_ receptor)^+^ GABAergic neurons suppressed the activity of surrounding LS cells, as concluded by decreased c-Fos-immunoreactivity [43]. Our data demonstrating both depolarization and hyperpolarization of dLS^GLP-1R^ neurons upon chemogenetic activation are consistent with this observation.

It is well established that the LS is strongly connected with the hypothalamus [44], which critically mediates feeding behavior and energy homeostasis [28]. Within the hypothalamus, the LHA also exerts control over motivated behavior, feeding, and energy balance across species [45]. Here, we found that dLS^GLP-1R^ neurons provide GABAergic input to the LHA. Our data demonstrate that chemogenetic or optogenetic stimulation of dLS^GLP-1R^→LHA neurons suppresses food intake, whereas chemogenetic inhibition of this pathway increase feeding. These results suggest a feeding-specific pathway from dLS to LHA. The LHA contains many molecularly and functionally distinct cell types [46–49]. Among these, GABAergic LHA neurons promote food seeking, consumption, and positive energy balance [45; 50]. We therefore speculate that GABAergic neurons are the downstream target of dLS^GLP-1R^ neurons within the LHA. Our data are consistent with this possibility because reduced inhibitory tone onto LHA GABA neurons (e.g., during chemogenetic inhibition of dLS^GLP-1R^ neurons) is expected to activate these cells and consequently increase feeding.

For decades GLP-1 has been known to reduce food intake by acting as a short-term prandial signal [6]. However, GLP-1 is also produced in the brain and is involved in a satiation/satiety circuit controlling food intake and body weight [35; 51; 52]. GLP-1 actions are mediated by GLP-1R, which is widely expressed in both the periphery [53] and the brain, including the dLS, LHA, paraventricular hypothalamic nucleus, among other regions. GLP-1R agonists, such as the analog of GLP-1 (liraglutide), have been used clinically as therapies to combat obesity [11]. In this study, we also find that Exn4, a specific GLP-1R agonist, can increase the optically induced IPSCs and decrease the presynaptic release probability at the dLS^GLP-1R^→LHA synapse, suggesting that GLP-1 signaling may mediate this pathway’s influence on feeding behavior.

In this study, we found that dLS^GLP-1R^ neurons and their inhibitory projections to the LHA can bidirectionally regulate food intake. These results extend our understanding of the neural mechanisms underlying feeding behavior and energy homeostasis. However, important unresolved questions remain. We have found local inhibitory connections within the dLS, which may have complex circuit architecture. For this reason, the specific mechanisms and functions of this local inhibition remain unclear. Moreover, while we speculate that the downstream target of dLS^GLP-1R^→LHA neurons are GABAergic neurons, further studies are required to elucidate these details.

## 5. Conclusions

Here, we dissected the role of dLS^GLP-1R^ neurons and their projections to the LHA in regulating feeding behavior. We showed that manipulations of both dLS^GLP-1R^ neurons or their projections to the LHA can influence feeding behavior independent of changes to anxiety-like behavior. A subpopulation of dLS^GLP-1R^ neurons send monosynaptic projections to the LHA where they release GABA, and the strength of this synapse is enhanced by GLP-1 signaling.

## CRediT authorship contribution statement

**Yi Lu:** Writing-original draft, Investigation. **Le Wang:** Investigation. **Fang Luo:** Investigation. **Rohan Savani:** investigation. **Mark Rossi:** Writing – review & editing, Writing – original draft, Investigation. **Zhiping Pang:** Writing – review & editing, Writing – original draft, Investigation.

## Acknowledgments

We thank the members of the Pang and Rossi laboratories for their suggestions and support for this study. We want to thank the Robert Wood Johnson Foundation for supporting the Child Health Institute of New Jersey (RWJF #74260).

## Data availability

Data will be made available on request.

## Figure legends

**Figure S1.**
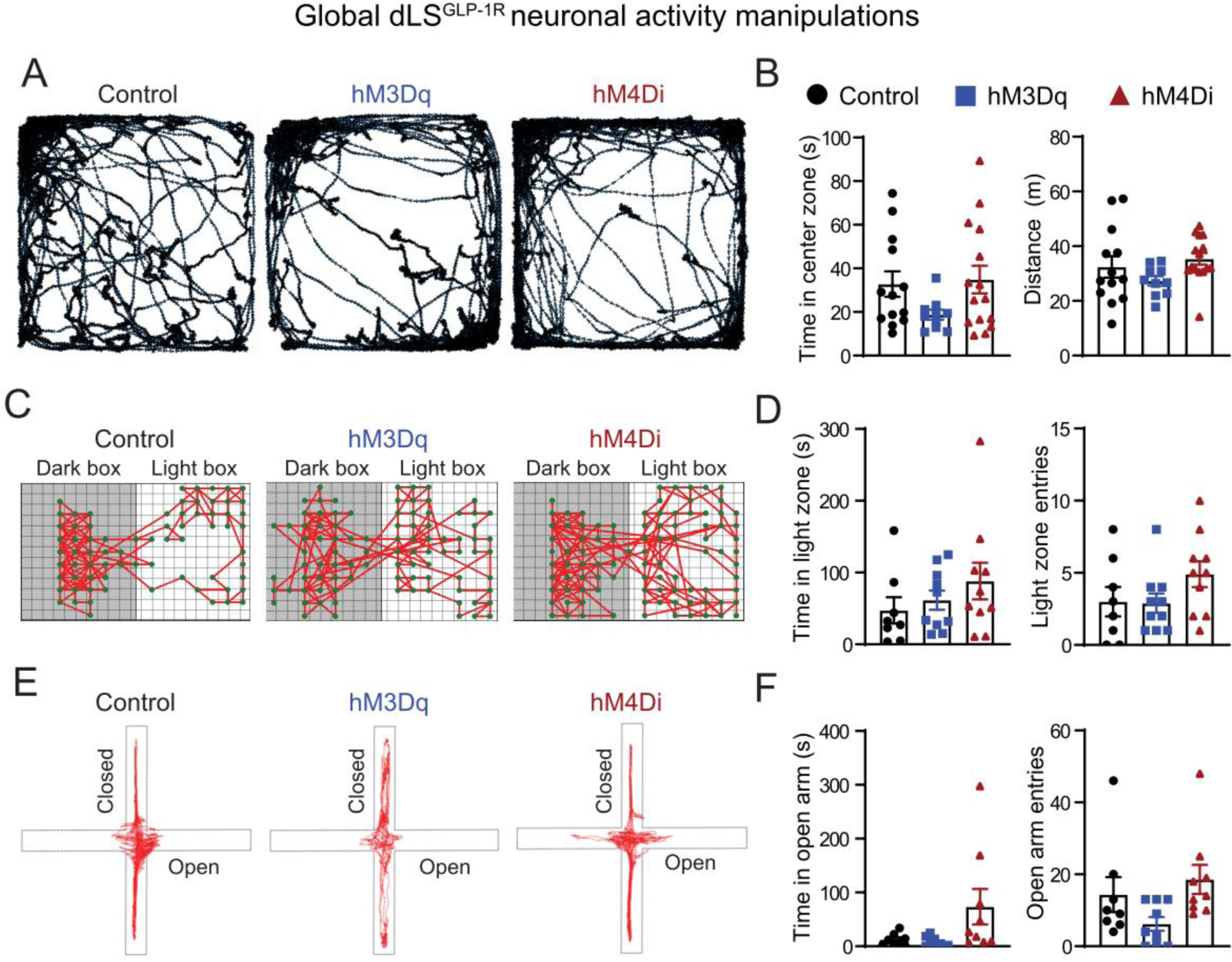
Chemogenetic manipulation of dLS^GLP-1R^ neurons does not affect anxiety-like behaviors. (A) Representative trajectories of AAV-DIO-mCherry-(left), AAV-DIO-hM3Dq-(middle) and AAV-DIO-hM4Di (right) injected mice during the open field test. (B) Quantification of time spent in the center zone (one-way ANOVA, F(2,35)=2.11, p=0.1364) and travel distance (one-way ANOVA, F(2,35)=1.898, p=0.1649) of open field test. No significant difference was detected between the groups. Data are presented as mean ±SEM. n=13 control, 10 hM3Dq, and 15 hM4Di mice. (C) Representative trajectories of AAV-DIO-mCherry-(left), AAV-DIO-hM3Dq-(middle) and AAV-DIO-hM4Di (right) injected mice during the light-dark box test. (D) Quantification of time spent in the light zone (one-way ANOVA, F(2,25)=1.041, p=0.3679) and entries into the light zone of light-dark box test (one-way ANOVA, F(2,25)=1.813, p=0.1840). No significant difference was detected between the groups. Data are presented as mean ± SEM. n=8 controls, 10 hM3Dq, and 10 hM4Di mice. (E) Representative trajectories of AAV-DIO-mCherry-(left), AAV-DIO-hM3Dq-(middle) and AAV-DIO-hM4Di (right) injected mice during the elevated plus maze test. (F) Quantification of time in open arm (one-way ANOVA, F(2,23)=3.414, p=0.0503) and open arm entries (one-way ANOVA, F(2,23)=2.941, p=0.0729) of elevated plus maze test. No significant difference was detected between the groups. Data are presented as mean ±SEM. n=8 controls, 9 hM3Dq, and 9 hM4Di mice.

**Figure S2.**
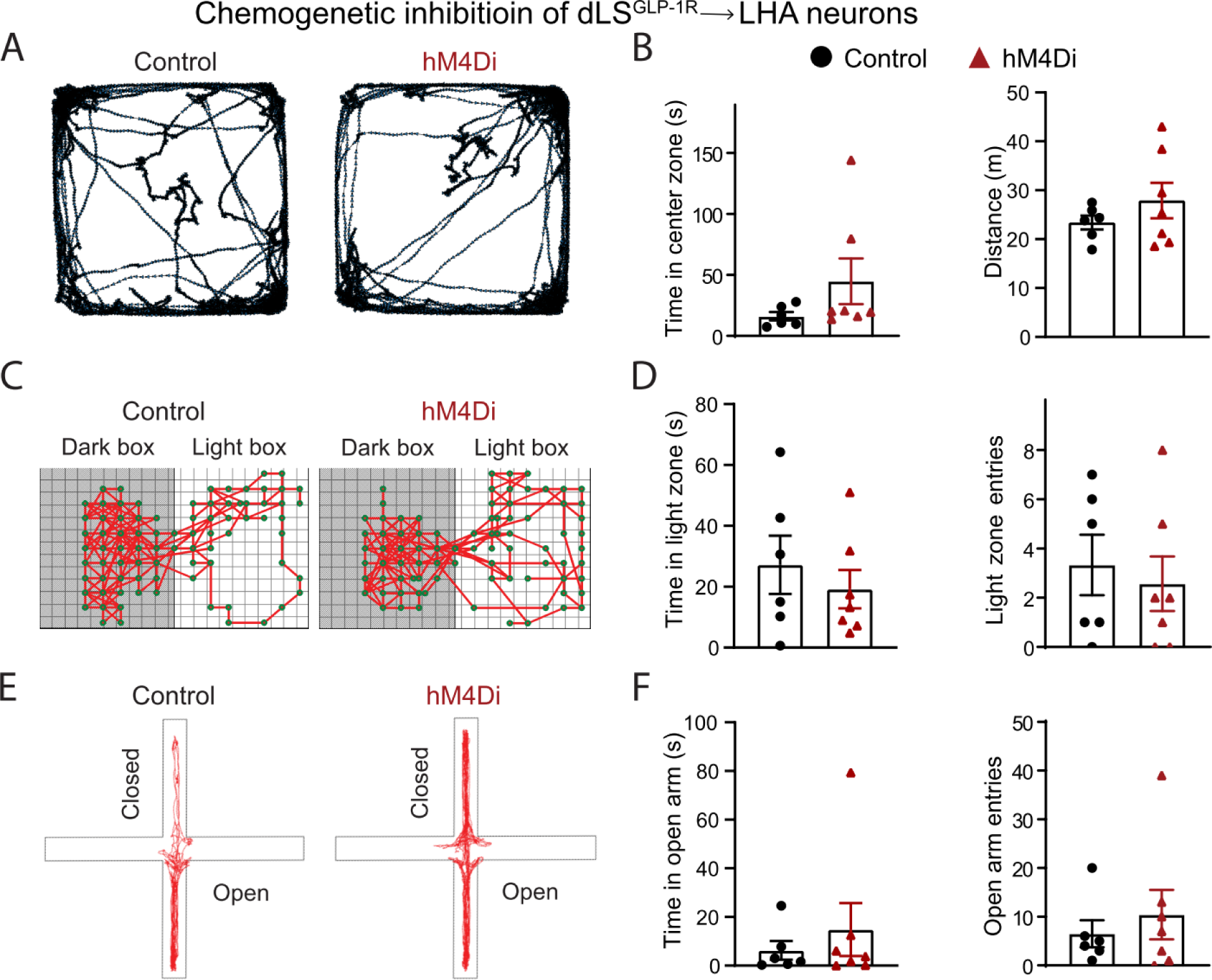
Chemogenetic inhibition of dLS^GLP-1R^→LHA projection neurons does not affect anxiety-like behaviors. (A) Representative trajectories of AAV-fDIO-mCherry-(left) and AAV-fDIO-hM4Di-injected (right) mice during the open field test. (B) Quantification of time in the center zone (unpaired t-test, t=1.397, p=0.1900) and travel distance of open field test (unpaired t-test, t=1.085, p=0.3012). No significant difference was detected between the groups. (C) Representative trajectories of AAV-fDIO-mCherry-(left) and AAV-fDIO-hM4Di-injected (right) mice during the light-dark box test. (D) Quantification of time spent in the light zone (unpaired t-test, t=0.7167, p=0.4885) and entries into the light zone of light-dark box test (unpaired t-test, t=0.4611, p=0.6537). No significant difference was detected between the groups. (E) Representative trajectories of AAV-fDIO-mCherry-(left) and AAV-fDIO-hM4Di-injected (right) mice during the elevated plus maze test. (F) Quantification of time spent in open arms (unpaired t-test, t=0.6903, p=0.5043) and open arm entries (unpaired t-test, t=0.6443, p=0.5326) of elevated plus maze test. No significant difference was detected between the groups. Data are presented as mean ±SEM, n = 6 control mice and 7 hM4Di mice.

**Figure S3.**
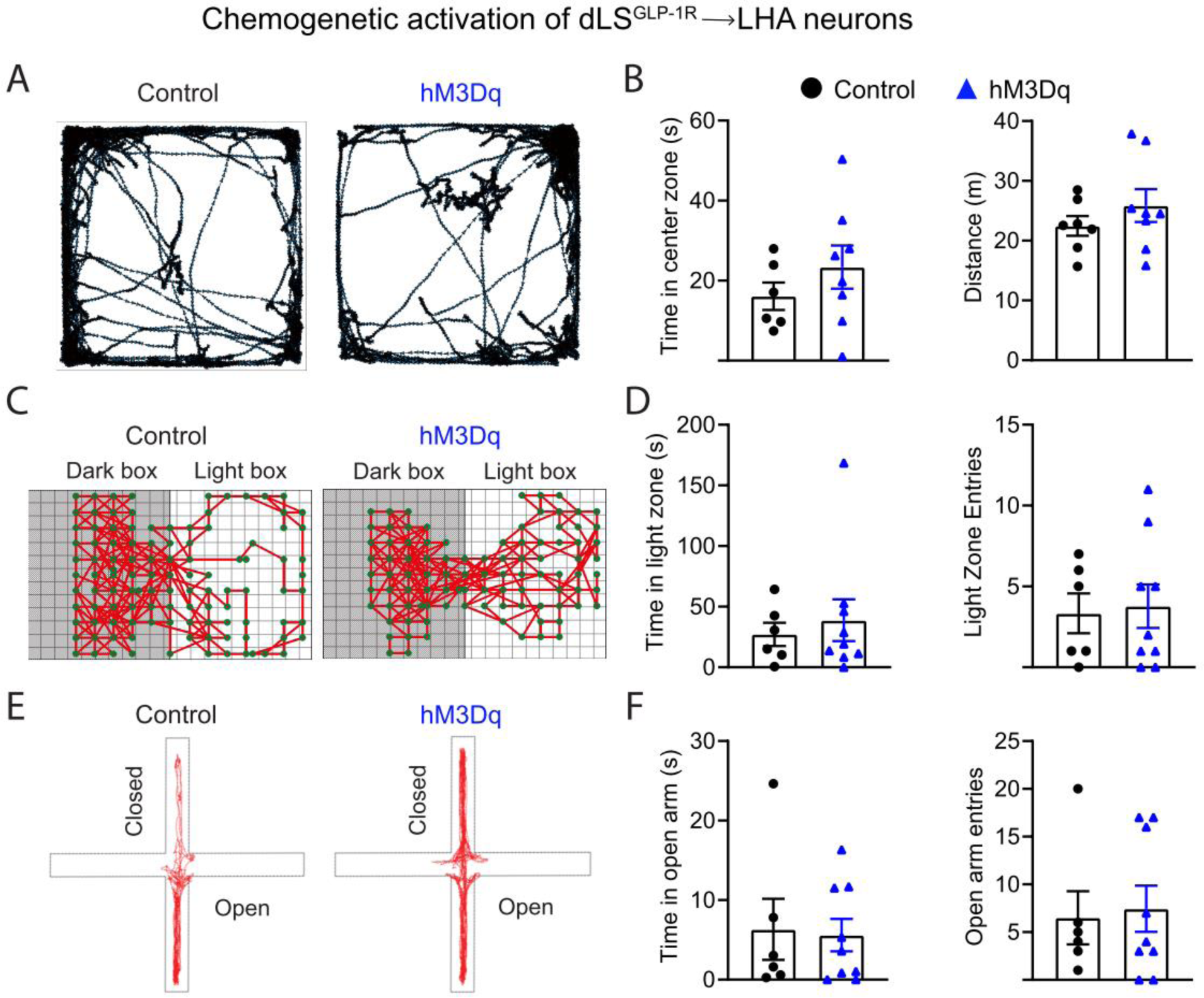
Chemogenetic activation of dLS^GLP-1R^→LHA projection neurons does not affect anxiety-like behaviors. (A) Representative trajectories of AAV-fDIO-mCherry-(left) and AAV-fDIO-hM3Dq-injected (right) mice during the open field test. (B) Quantification of time in the center zone (unpaired t-test, t=1.044, p=0.3171) and travel distance of open field test (unpaired t-test, t=0.7180, p=0.4865). No significant difference was detected between the groups. Data are presented as mean ±SEM, n=6 control mice and 8 hM3Dq mice. (C) Representative trajectories of AAV-fDIO-mCherry-(left) and AAV-fDIO-hM3Dq-injected (right) mice during the light-dark box test. (D) Quantification of time spent in the light zone (unpaired t-test, t=0.5106, p=0.6182) and entries into the light zone of light-dark box test (unpaired t-test, t=0.2299, p=0.8217). No significant difference was detected between the groups. Data are presented as mean ±SEM, n=6 control mice and 9 hM3Dq mice. (E) Representative trajectories of AAV-fDIO-mCherry-(left) and AAV-fDIO-hM3Dq-injected (right) mice during the elevated plus maze test. (F) Quantification of time spent in open arms (unpaired t-test, t=0.1821, p=0.8583) and open arm entries (unpaired t-test, t=0.2531, p=0.8042) of elevated plus maze test. No significant difference was detected between the groups. Data are presented as mean ±SEM, n=6 control mice and 9 hM3Dq mice.

